# Analysis of nucleic acids extracted from rapid diagnostic tests reveals a significant proportion of false positive test results associated with recent malaria treatment

**DOI:** 10.1101/2021.05.12.443950

**Authors:** Salome Hosch, Charlene Aya Yoboue, Olivier Tresor Donfack, Etienne A. Guirou, Jean-Pierre Dangy, Maxmillian Mpina, Elizabeth Nyakurungu, Koranan Blöchliger, Carlos A. Guerra, Wonder P. Phiri, Mitoha Ondo’o Ayekaba, Guillermo A. García, Marcel Tanner, Claudia Daubenberger, Tobias Schindler

## Abstract

Surveillance programs often use malaria rapid diagnostic tests (RDTs) to determine the proportion of the population carrying parasites in their peripheral blood to assess the malaria transmission intensity. Despite an increasing number of reports on false-negative and false-positive RDT results, there is a lack of systematic quality control activities for RDTs deployed in malaria surveillance programs. Our study provides a larger scale comparative evaluation of RDTs used in the 2018 Malaria Indicator Survey (MIS) conducted on Bioko Island, Equatorial Guinea. We conducted a molecular analysis by extraction of nucleic acids from 1,800 negative and 1,065 positive RDTs followed by qPCR analysis. These results were combined with a dataset collected in a comprehensive questionnaire from each MIS participant. Of the 2,865 RDTs that were collected in 2018 on Bioko Island and analysed in our study, 4.7% had a false-negative result. These false-negative RDT results were associated with low parasite density infections. In a substantial proportion of samples, we identified masked *pfhrp2* and *pfhrp3* gene deletions in which at least one *P. falciparum* strain carried a gene deletion. Among all positive RDTs analysed, 28.4% were tested negative by qPCR and therefore considered to be false-positive. Analysing the questionnaire data collected from the participants, this high proportion of false-positive RDT results could be explained by PfHRP2 antigen persistence after recent malaria treatment. We conclude that malaria surveillance depending solely on RDTs needs well-integrated quality control procedures assessing the extend and impact of reduced sensitivity and specificity of RDTs on malaria control programs.

## Introduction

According to the World Health Organization, more than 409,000 malaria deaths were reported in 2019, most of them in children below the age of five years ^1^. The majority of malaria infections (94%) and malaria related deaths (95%) occurred in sub-Saharan Africa (sSA), where *Plasmodium falciparum (Pf)* is the dominant malaria parasite ^1^. The test-treat-track strategy advised by WHO is one of the backbones of current malaria control and elimination programs ^2^. This strategy entails that every suspected malaria case should be tested, every confirmed case should be treated, and the disease should be tracked through surveillance systems ^3^. Testing relies heavily on rapid diagnostic tests (RDTs), exemplified by the more than 348 million RDTs distributed globally in 2019 ^1^. In sSA, RDTs have almost completely replaced light microscopy for malaria diagnosis, accounting for an estimated 75% of all malaria tests conducted in 2017 ^4^. RDTs are point-of-care tests that detect circulating antigens like the *Pf* specific histidine rich protein 2 (PfHRP2) or histidine rich protein 3 (PfHRP3) and the pan-*Plasmodium* spp. enzymes lactate dehydrogenase (pLDH) or aldolase ^5^. More than 90% of RDTs currently in use target the PfHRP2 antigen because of its higher sensitivity compared to non-PfHRP2 antigens ^6^. PfHRP2-based RDTs used for the diagnosis of febrile patients that suffer from malaria infection are highly sensitive and specific ^7^. RDTs are often used by national malaria surveillance programs. However, when individuals are asymptomatic with low parasite densities, RDTs often fail to detect the parasites due to low antigen concentrations ^8, 9^. A recent study showed that false-negative RDTs (FN-RDT) are more common in lower malaria transmission settings, younger subjects, and in urban areas in sSA ^10^. Reduced diagnostic performance of RDTs has also been attributed to genetic diversity of the *pfhrp2* gene ^11^, differences in expression levels of PfHRP2 antigen in parasite field strains ^12^, or deletion of *pfhrp2* and *pfhrp3* genes in isolates ^13^. *Pfhrp2* gene deletions appear to be common and therefore are relevant as they might be a threat to malaria control programs based on monitoring of malaria prevalence through RDT ^14, 15^.

Less attention has been given to the specificity of malaria RDTs used in malaria surveys that potentially result in false positive results. False-positive RDTs (FP-RDT) have been associated with high levels of circulating rheumatoid factor ^16–18^ or acute typhoid fever ^19^. There is evidence of FP-RDTs in patients infected with *Schistosoma mekongi* ^20^ or human African trypanosomiasis ^21^. FP-RDTs are also caused by persisting antigen circulation in peripheral blood after malaria drug treatment. A meta-analysis revealed that half of the PfHRP2-detecting RDTs remain positive 15 (95% CI: 5-32) days post malaria treatment, 13 days longer than RDTs targeting the pLDH antigen ^22^. The latter study also reported a higher persistent RDT positivity among individuals treated with artemisinin combination therapy (ACT) than those treated with other anti-malarial drugs. RDTs are instrumental to malaria surveillance programs, and therefore their diagnostic performance should be systematically monitored over time using molecular methods detecting *Plasmodium* spp. genomic markers. We describe here an approach for quality control of field-deployed malaria RDTs by retrospective molecular analysis of the parasite DNA retained on them using qPCR.

## Materials and Methods

### The 2018 malaria indicator survey conducted on Bioko Island as a biobank of RDTs for molecular malaria surveillance

A malaria indicator survey has been conducted annually since 2004 on the Island of Bioko, Equatorial Guinea, to evaluate the impact of malaria control interventions ^23^. The survey uses a standard questionnaire developed by the Roll Back Malaria initiative to gather information on selected households and their occupants. The 2018 Bioko Island MIS covered 4,774 households with 20,012 permanent residents, among whom 13,505 persons consented to storage and molecular analysis of their RDT. Briefly, consenting individuals living in surveyed households are tested for malaria and malaria-related anaemia. Malaria testing was done with the CareStart^™^ Malaria HRP2/pLDH (Pf/PAN) combo test (ACCESS BIO, New Jersey, USA). The haemoglobin level in peripheral blood was measured during the MIS using a battery-operated portable HemoCue system (HemoCue AB, Ängelholm, Sweden). The anaemia status (mild, moderate, severe) was categorized based on definitions published by the World Health Organization ^24^ stratified by age, gender, and pregnancy status. Households were assigned scores based on the type of assets and amenities they own to derive a surrogate of their socio-economic status (SES), using principal component analysis (PCA). After ranking all households based on their score, they were divided into five equal categories (quintiles), each with approximately 20% of the households. The first quintile corresponded to the lowest wealth index and the fifth to the highest wealth index. The household wealth index categories were also assigned to permanent household members.

### Detection and quantification of *Plasmodium* spp. nucleic acids extracted from RDTs

We developed an approach to extract nucleic acids (NA) from blood retained on used malaria RDTs named “**E**xtraction of **N**ucleic **A**cids from **R**DTs” (ENAR) ^25^. Briefly, RDTs were barcoded, stored at room temperature, and shipped to Basel, Switzerland, for NA extraction and detection. This approach simplifies small volume blood collection, transport and storage logistics, and allows linking outcomes of molecular based detection of parasite derived NA with the demographic and socio-economic information collected from each corresponding MIS participant at high throughput.

In total, 2,865 RDTs (21.2%) collected during the 2018 MIS were included in this study. The median age in this sample collection was 22 years (interquartile range 9 to 38 years), female participants were slightly overrepresented (58.2%), and 97.8% of the participants were asymptomatic, non-febrile individuals. More than two-thirds of the RDTs were collected in the urban areas of the capital city Malabo on Bioko Island.

All 2,865 samples were initially screened with the PlasQ RT-qPCR assay ^26^. In this RT-qPCR assay, the high copy number *Pf* specific varATS region ^27^ and the pan-*Plasmodium* 18S rDNA gene were targeted ^28, 29^. Samples with cycle of quantification (Cq) value < 45 in two replicates of either of the two targets, varATS or 18S rDNA, were considered positive. *Pf* parasites were quantified based on their Cq value for varATS ^25^. In addition, only samples with Cq value < 35 for amplification of the internal control gene, the human *rnasep* gene were included, to demonstrate that the NA extracted from the RDTs is sufficient for reliable molecular analysis of malaria parasites. Non-*falciparum* malaria species identification of samples positive for the pan-*Plasmodium* target 18S rDNA was performed with a multiplex RT-qPCR assay based on species-specific 18S rDNA sequences as described previously ^25^.

### Quality control and categorization of RDT outcomes

A RDT was considered positive if the healthcare worker recorded a positive signal for the PfHRP2, pLDH, or both targets during the MIS. Among these positive RDTs, a true-positive RDT (TP-RDT) result was defined as a RDT with detectable *Plasmodium* spp. NA (two replicates with varATS and/or 18S rDNA Cq < 45 and human *rnasep* Cq < 35). A false-positive RDT (FP-RDT) result was defined as positively read and recorded RDT in the field but with a negative outcome for *Plasmodium* spp. NA based on PlasQ RT-qPCR in the presence of human *rnasep* Cq < 35. Negative RDTs were classified as being read as negative by the healthcare worker during the MIS and recorded in the database. A true-negative RDT (TN-RDT) result was defined as a RDT whose negative result collected in the field was confirmed by the PlasQ RT-qPCR. A false-negative RDT (FN-RDT) result was defined as negatively read by the healthcare worker in the field with a positive PlasQ RT-qPCR result based on two replicate amplifications with varATS and/or 18S rDNA Cq < 45 and the human *rnasep* Cq < 35.

### qHRP2/3-del assay for detection of *pfhrp2* and *pfhrp3* deletions

The previously published qHRP2/3-del assay that simultaneously amplifies the *pfhrp2* and *pfhrp3* genes together with the internal control gene *pfrnr2e2* was adapted to accommodate for the lower input of NA ^30^. Briefly, the probe for the internal control gene *pfrnr2e2* was labelled with fluorescein (FAM) instead of Cy5 to improve its detectability. Additionally, the final concentration of all primers was increased from 0.3 μM to 0.45 μM. Concentrations of 0.15 μM were used for the *pfrnr2e2* probe, and 0.225 μM for the *pfhrp2* and *pfhrp3* probes each. All samples were run in triplicates and the number of amplification cycles was increased from 45 to 50. Every 96 well qPCR plate contained control DNA extracted from a known *pfhrp2*-deleted Pf strain (Dd2), a *pfhrp3*-deleted Pf strain (HB3), and a Pf strain without *pfhrp2* and *pfhrp3* gene deletions (NF54) as well as a non-template control (NTC). We defined successful amplification as a mean Cq < 40 for *pfrnr2e2* calculated from at least two replicates for each sample. We ran the qHRP2/3-del assay only with NA extracted from RDTs that had displayed a Cq < 35 for the *varATS* target in the PlasQ RT-qPCR.

*Pfrnr2e2*, *pfhrp2,* and *pfhrp3* are all single-copy genes and they show comparable performances in the multiplex qPCR assay ^30^. One approach to detect Pf strains with *pfhrp2* and/or *pfhrp3* gene deletions in mixed Pf strain infections (herein defined as masked gene deletions) is to calculate the difference in Cq values obtained between *pfhrp2* or *pfhrp3* and *pfrnr2e2* amplifications (ΔCq values). This is done by subtracting the Cq value obtained during the amplification of *pfrnr2e2* from the Cq value of *pfhrp2* or *pfhrp3*, respectively. Combining all runs that were conducted, the mean ΔCq for *pfhrp2* in controls (NF54 and HB3) was 0.00 (SD±0.52) and for *pfhrp3* the mean ΔCq in controls (NF54 and Dd2) was 1.19 (SD ±0.83). For *pfhrp2* a ΔCq cut-off value of 1.0 (mean + 2x SD for controls) was chosen to identify masked gene deletions. For *pfhrp3* a ΔCq cut-off value of 2.9 (mean + 2x SD for controls) was chosen to identify masked gene deletions.

### Genotyping of *P. falciparum pfmsp1* and *pfmsp2 genes*

Genotyping with *pfmsp1* and *pfmsp2* was performed following published procedures using nested PCR ^31^. The first two PCR reactions amplify conserved sequences within the polymorphic regions of *pfmsp1* and *pfmsp2*, respectively. The second, nested PCR targets allele-specific sequences in five separate reactions. Samples were run in 20 μL total volume with 1x Hot Firepol Master Mix (Solys BioDyne, Estonia), 0.25 μM of forward and reverse primers and 2 μL template DNA. The cycling conditions for the first PCR were 95°C for 12 minutes, 25 cycles of 95°C for 30 seconds, 58°C for 1 minute and 72°C for 2 minutes and 72°C for 10 minutes. For the second PCR, the cycling conditions for the three allele-specific *pfmsp1* primer pairs were 95°C for 12 minutes, 35 cycles of 95°C for 30 seconds, 56°C for 40 seconds and 72°C for 40 seconds and 72°C for 10 minutes. For the two *pfmsp2* allele-specific reactions the conditions were: 95°C for 12 minutes, 35 cycles of 95°C for 30 seconds, 58°C for 40 seconds and 72°C for 40 seconds and 72°C for 10 minutes. Presence and size of PCR products was determined and documented visually on a 1% agarose gel with a 100bp DNA ladder.

### Genotyping of *P. malariae* circumsporozoite protein (pmcsp)

The *pmcsp* gene was amplified by semi-nested PCR for all samples with a positive signal for *P. malariae* in the non-*falciparum* malaria species identification assay ^25^. The first PCR was run with 3 μL of DNA template in a reaction volume of 20 μL. The reaction mix contained 1x Hot Firepol Master Mix and 0.25 μM of each of the primers csp_OF ^32^ and csp-R ^33^. The conditions for the first PCR were: 95 °C for 12 minutes; 35 cycles of 95 °C for 15 seconds, 53 °C for 30 seconds and 65 °C for 90 seconds and final elongation at 65 °C for 10 minutes. The second, semi-nested PCR used 1.5 μL of the product from the first reaction in a total volume of 15 μL. The reaction mix contained 1x Hot Firepol Master Mix and 0.33 μM of the primers csp_IF ^32^ and csp-R. The conditions for the second PCR were: 95 °C for 12 minutes; 35 cycles of 95 °C for 15 seconds, 52 °C for 30 seconds and 62 °C for 90 seconds and final elongation at 62 °C for 10 minutes. The PCR product was sent to Microsynth (Microsynth AG, Switzerland) for bidirectional sanger sequencing.

### Data analysis and statistics

The generated (RT)-qPCR data was initially analysed with the CFX Maestro Software (Bio-Rad Laboratories, California, USA). Thresholds for each fluorescence channel were set manually and Cq values were then uploaded to the ELIMU-MDx platform for data storage and analysis (23). Sequence analysis was performed using Geneious Prime 2019.1.1 (https://www.geneious.com). Statistical analysis and data visualisation was performed using the R statistical language (version 4.0.3) based on packages data.table, dplyr, epiDisplay, epitools, ggplot2, ggpubr, ggridges, gridExtra, lme4, readxl, reshape2, scales, stringr, tidyr, tidyverse. Wilcoxon rank sum test was used for numeric values. Fisher’s exact test (two-sided) was used for contingency tables. A generalized linear mixed-effects model with fixed and random effects was used for calculation of odds ratios and their confidence intervals.

### Data availability

All data needed to evaluate the conclusions in the paper are present in the manuscript or the Supplementary Materials. Further information will be made available to interested researchers. The 15 sequences of *P. malariae* circumsporozoite protein from Bioko Island have been deposited into GenBank under the accession numbers MW963324-MW963338.

### Ethical approval

The Ministry of Health and Social Welfare of Equatorial Guinea and the Ethics Committee of the London School of Hygiene & Tropical Medicine (Ref. No. LSHTM: 5556) approved the 2018 malaria indicator survey. Written informed consent was obtained from all adults and from parents or guardians of children who agreed to participate. Only samples for which an additional consent for molecular analysis was obtained were included in this study. We confirm that all experiments were performed in accordance with relevant national and international guidelines and regulations.

## Results

### Integration of molecular diagnostic methods into the national malaria control program to assess the performance of malaria RDTs

Following NA extraction, a PlasQ RT-qPCR result was generated for 1,800 malaria negative and 1,065 malaria positive RDTs, as collected in the MIS database. By comparison between PlasQ RT-qPCR results and RDT results collected in the field, RDTs were grouped into four categories, namely true-positive (TP), true-negative (TN), false-positive (FP), and false-negative (FN), respectively (Figure 1). The PlasQ RT-qPCR was used as a gold standard to evaluate the performance of the RDT, and this resulted in an overall sensitivity of 90.0% and specificity of 85.0% of field-deployed RDTs used during the 2018 Bioko Island MIS.

**Figure 1.**
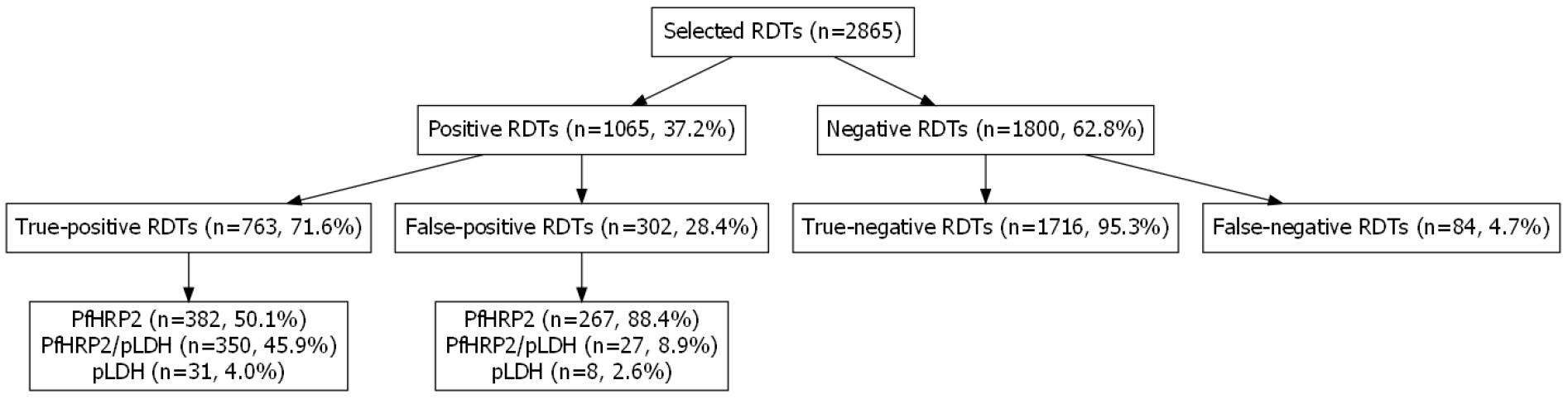
Comparison of RDT outcomes collected during 2018 MIS with PlasQ RT-qPCR results obtained after NA extraction and amplification.

When stratified by the type of antigen, FP-RDTs were predominantly those that detected only the PfHRP2 antigen (88.4%) during the MIS; whereas 8.9% and 2.6% of the FP-RDTs were respectively those that detected both, the PfHRP2 plus the pLDH antigens, and those that detected the pLDH antigen only. Around half of TP-RDTs were those that detected the PfHRP2 antigen only (50.1%), followed by those that detected both antigens (45.9%) and lastly, those that detected the pLDH antigen only (4.0%) during the MIS.

### Low parasite density infections are likely to cause false-negative RDT results in the field

The ENAR approach used in this study detects 10-100x lower parasite densities than the PfHRP2-based RDT itself ^25^. Here, we confirm that a clear association exists between FN-RDT, TP-RDT, and *Pf* parasite densities assessed by the PlasQ RT-qPCR outcome. TP-RDT had higher geometric mean parasite densities (35.0 Pf/μL, IQR: 7.2-166.0) compared to FN-RDTs (4.6 Pf/μL, IQR: 1.1-20.0) (Figure 2a, Wilcoxon rank sum test, p<0.001). Although *Pf* was the most common (93.8%) *Plasmodium* spp. species among RT-qPCR positive RDTs, *P. malariae* (4.0%) and *P. ovale* spp. (1.1%) were also identified. No *P. vivax* and *P. knowlesi* parasite NAs were detected. The central repeat region of the *P. malariae* circumsporozoite protein (*pmcsp*) was amplified by PCR and Sanger sequenced to confirm the presence of *P. malariae* (Supplementary figure 1b). Nucleotide sequences were unique among all 15 *P. malariae* isolates sequenced and also the number of NAAG and NDAG repeats varied between the isolates. These results indicated high diversity of the *P. malariae* population on Bioko Island. A slightly higher proportion, although not statistically significant, of non-*falciparum Plasmodium* spp. parasites was found among FN-RDTs. *P. malariae* was found among 6.6% of FN-RDTs compared to 3.8% among TP-RDTs. Similar, *P. ovale* spp. was more prevalent in FN-RDTs (2.6%) than in TP-RDTs (0.9%).

**Figure 2.**
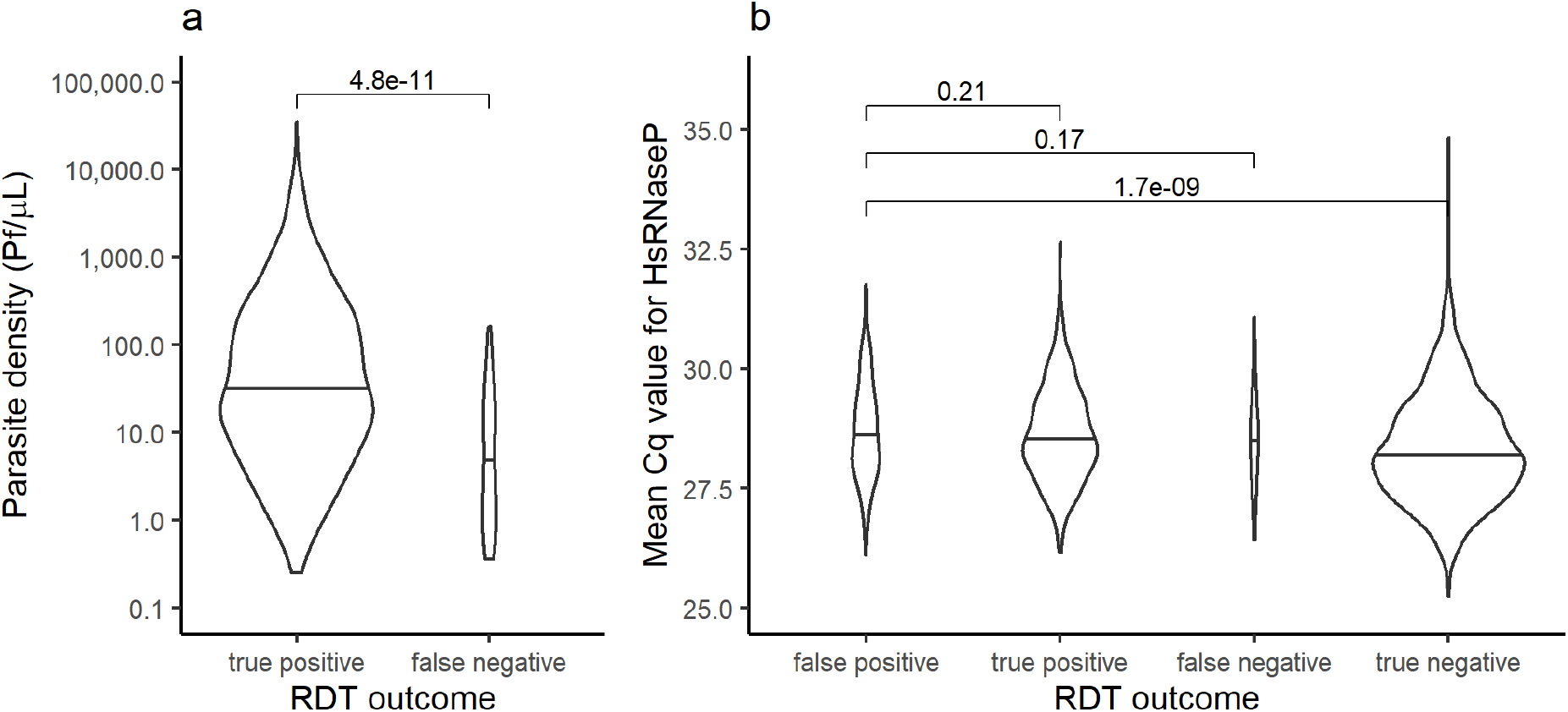
(a) *Pf* parasite densities compared between true positive and false negative RDT outcomes. Parasite densities for *Pf* were estimated based on the varATS target of the PlasQ RT-qPCR assay. Wilcoxon rank sum test was used to compare the two groups. (b) Comparison of the Cq values obtained with the amplification of the human *rnasep* gene used as internal control of the PlasQ RT-qPCR assay, across all samples stratified by RDT outcome. The group of RDTs with a false-positive result was compared to the other RDT outcomes by Wilcoxon rank sum test.

### False-negative RDT results are not associated with parasites carrying *pfhrp2* and *pfhrp3* gene deletions

*Pf* strains were genotyped to identify *pfhrp2* and/or *pfhrp3* gene deletions. The number of samples available was limited through the combination of low parasite density infections and the limited amount of blood retained on RDTs as a source of NA. The single copy gene *pfrnr2e2*, serving as the internal control of the qHRP2/3-del assay, was amplified with Cq < 40 in 184/406 (45.3%) samples. To avoid false reporting of *pfhrp2* and/or *pfhrp3* gene deletions, the analysis was restricted to samples that had an additional amplification in either *pfmsp1* (32/47, 68.1%) or *pfmsp2* (31/47, 66.0%). The success rate as a function of the parasite density for amplifying each genotyping marker based on parasite density is shown in Figure 3a-c. At least two out of three reference genes (*pfrnr2e2, pfmsp1* or *pfmsp2*) were amplified in thirty-six samples, which were then included in the analysis of the *pfhrp2* and *pfhrp3* deletion status. No evidence for parasites carrying a *pfhrp2* gene deletion was found in these 36 samples, but four out of 36 samples (11.1%) were likely to carry *pfhrp3* gene deletions. All four samples with *pfhrp3* deletion were recorded as positive for PfHRP2 by RDT.

**Figure 3.**
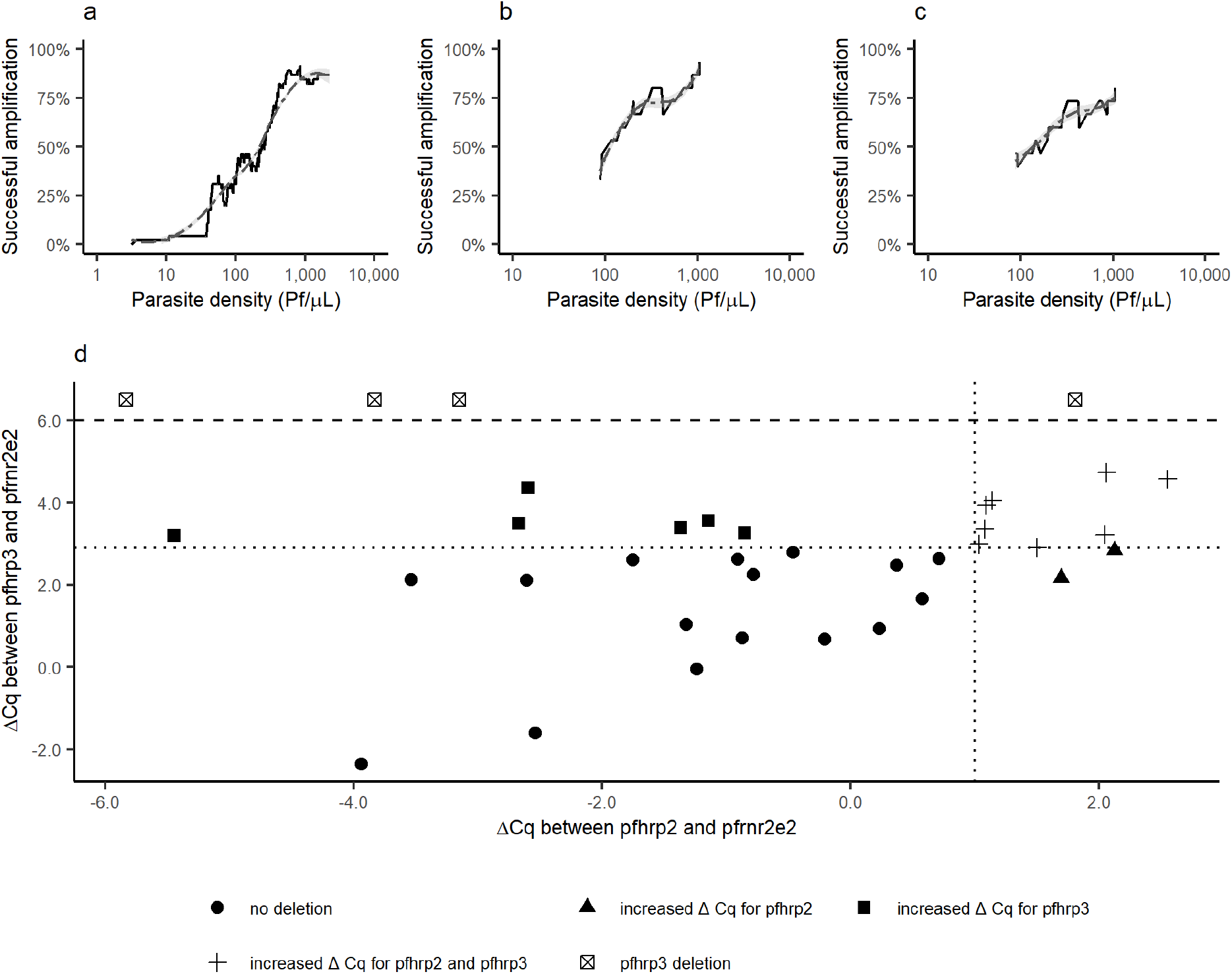
Amplification rate (rolling mean) for the genotyping reference genes (a) *pfrnr2e2*, (b) *pfmsp1* and (c) *pfmsp2* as a function of the parasite density of the sample. Parasite densities for *Pf* were estimated based on the *varATS* target of the PlasQ assay. (d) The distribution of ΔCq values between *pfhrp2* (x-axis) or *pfhrp3* (y-axis) and *pfrnr2e2*. ΔCq thresholds (dashed lines) were set at 1.0 for *pfhrp2* and 2.9 for *pfhrp3*. For samples with a *pfhrp3* deletion, the ΔCq for *pfhrp3* was set arbitrarily at 6.5.

Based on the available data from the *pfmsp1* and *pfmsp2* genotyping (Supplementary figure 1a), polyclonal infections consisting of two or more distinct *Pf* clones were found in 63.0% (17/27) of samples with successful amplification of *pfmsp1* and *pfmsp2*. The qHRP2/3-del assay was used to identify *pfhrp2* and/or *pfhrp3* gene deletions in polyclonal *Pf* infections by calculating the ΔCq values as the difference of Cq values between *pfhrp2 and pfhrp3* gene amplification and the *pfrnr2e2* internal control. Figure 3d shows the distribution of samples with their respective ΔCq values for *pfhrp2* and *pfhrp3.* Of the 36 samples included, nine samples (25.0%) had increased ΔCq values for both genes, two samples (5.6%) only for the *pfhrp2* gene and six samples (16.7%) only for the *pfhrp3* gene, respectively. Importantly, the 11 samples which had an increased ΔCq for the *pfhrp2* gene were positive for PfHRP2 by RDT.

### False-positive RDT results are associated with recent use of antimalarial drugs

The rate of FP-RDTs differed across age, level of anaemia, geographical location of residence, and the socio-economic status (Supplementary figure 2). Interestingly, no study participant with a FP-RDT had a fever (>37.5° C) at the time of the survey, while 1.7% (13/763) of those with TP-RDTs were recorded with fever. Eight factors from the MIS were used to identify risk factors associated with FP-RDTs through multivariate logistic regression analysis in which the outcome of the test was set as the outcome variable (Supplementary table 1). FP-RDTs (n=297) were compared to TP-RDTs (n=754). Because sample collection was clustered within communities, community affiliation was introduced as a random effect to the model. The MIS included 299 communities, of which 201 (67.2%) were represented in the dataset. The median number of samples from a community was three. Survey participants belonging to higher socio-economic classes (aOR 1.51 p=0.01) had increased odds of having a FP-RDT. Participants who were reported to have been treated with an antimalarial drug in the two weeks preceding the survey had more than four times the odds of a FP-RDT result than a TP-RDT (aOR 4.52, p<0.001). In contrast, moderate to severe anaemia reduced the odds of having a FP-RDT (aOR 0.60, p=0.02). Those who reside in the rural Bioko Sur province had also decreased odds of having a FP-RDT (aOR 0.44, p=0.01). Age, sex, bednet use, and reported sickness in the two weeks preceding the survey were not significantly associated with FP-RDTs (Figure 4).

**Figure 4.**
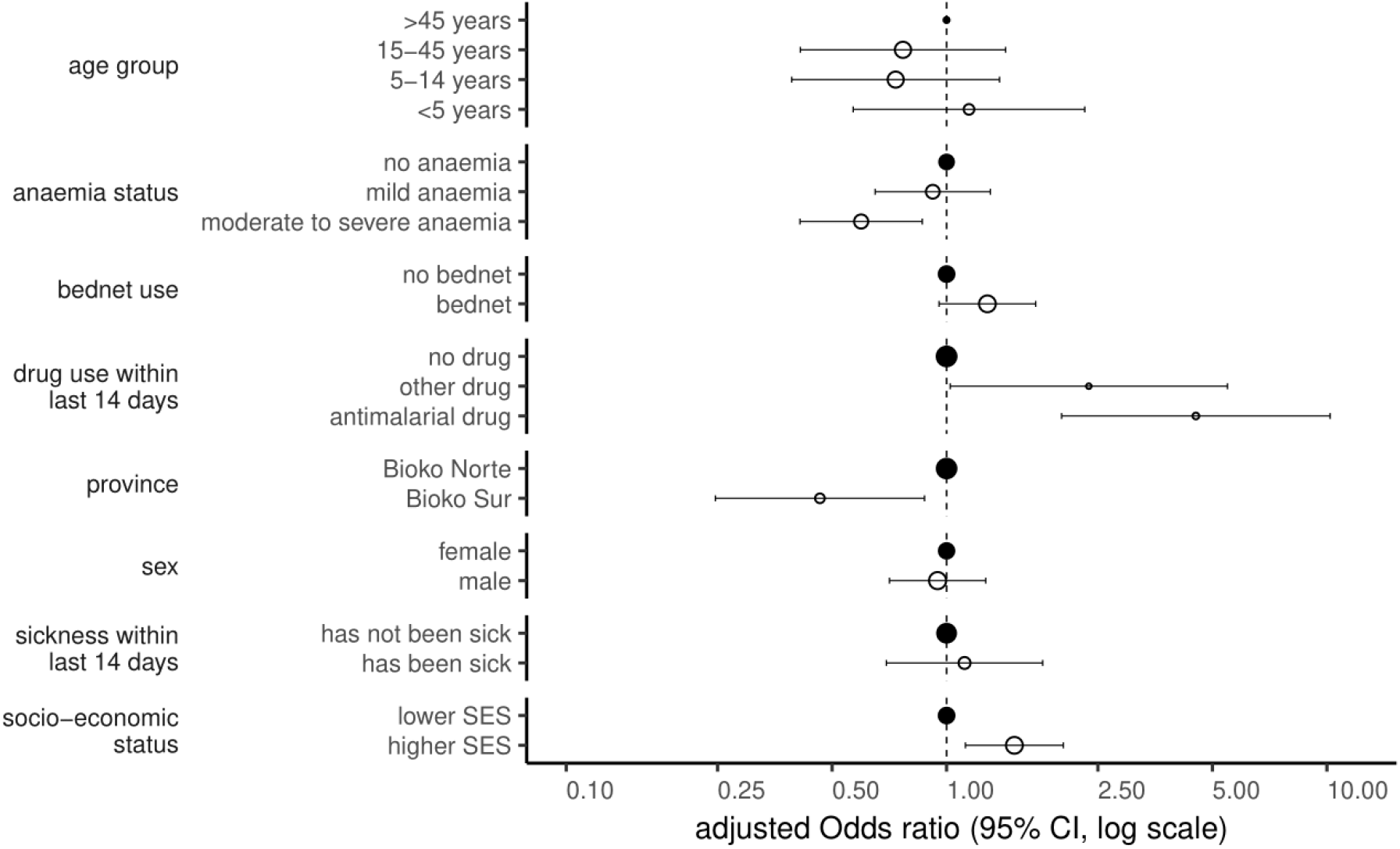
Risk factors associated with FP-RDT results by multivariate logistic regression analysis. The size of the circles corresponds with the number of responses for each variable outcome. The reference group is marked by filled circles and the other groups have open circles. Higher socio-economic status (SES) included people from the 4^th^ and 5^th^ wealth quintiles.

To exclude the possibility that FP-RDTs are the consequence of failed amplification related to the degradation of NA retained on the RDTs, an additional analysis was carried out. During the PlasQ RT-qPCR, the human *rnasep* gene was used as an internal control to monitor the amount of NA extracted from each RDT. On average, the human *rnasep* was amplified with a Cq value of 28.5 (SD ±1.0). There was no significant difference in the Cq values of the human *rnasep* gene amplification among RDTs, which were categorized as false positive (28.6, SD± 1.0), true positive (28.5, SD± 1.0), or false negative (28.6, SD± 1.0). TN-RDTs had a significantly lower median Cq value (28.2, SD± 1.1) (Figure 2b). These results indicate that the lack of detectable *Pf* NA in the blood retained on FP-RDTs is not related to poor NA extraction performance or a failure in detecting NAs.

### The impact of asymptomatic malaria infections on anaemia status might be underestimated by false-positive RDT results

We hypothesised that high rates of FP-RDTs are likely to lead to underestimating the impact of asymptomatic malaria infections on anaemia status. Among malaria infected children aged <5 years, the prevalence of anemia was 67.7% if malaria status was assessed by RDT. Stratification by RT-qPCR-based correction revealed that the proportion of anemic children with a FP-RDT result (48.9%) is similar to children with a TN-RDT result (41.4%), whereas children with a TP-RDT result are more likely to suffer from anemia (78.3%) (Figure 5). This significant effect is even more pronounced among children with moderate and severe anemia if compared to mild anemia. Removing all FP-RDTs in this association between malaria infection status and anemia levels in children < 5 years reveals that the association between asymptomatic malaria and moderate or severe anaemia might be even higher than if malaria infection status is assessed by RDT only. In older children and adults, the impact of FP-RDTs on assessing the anaemia status is negligible.

**Figure 5.**
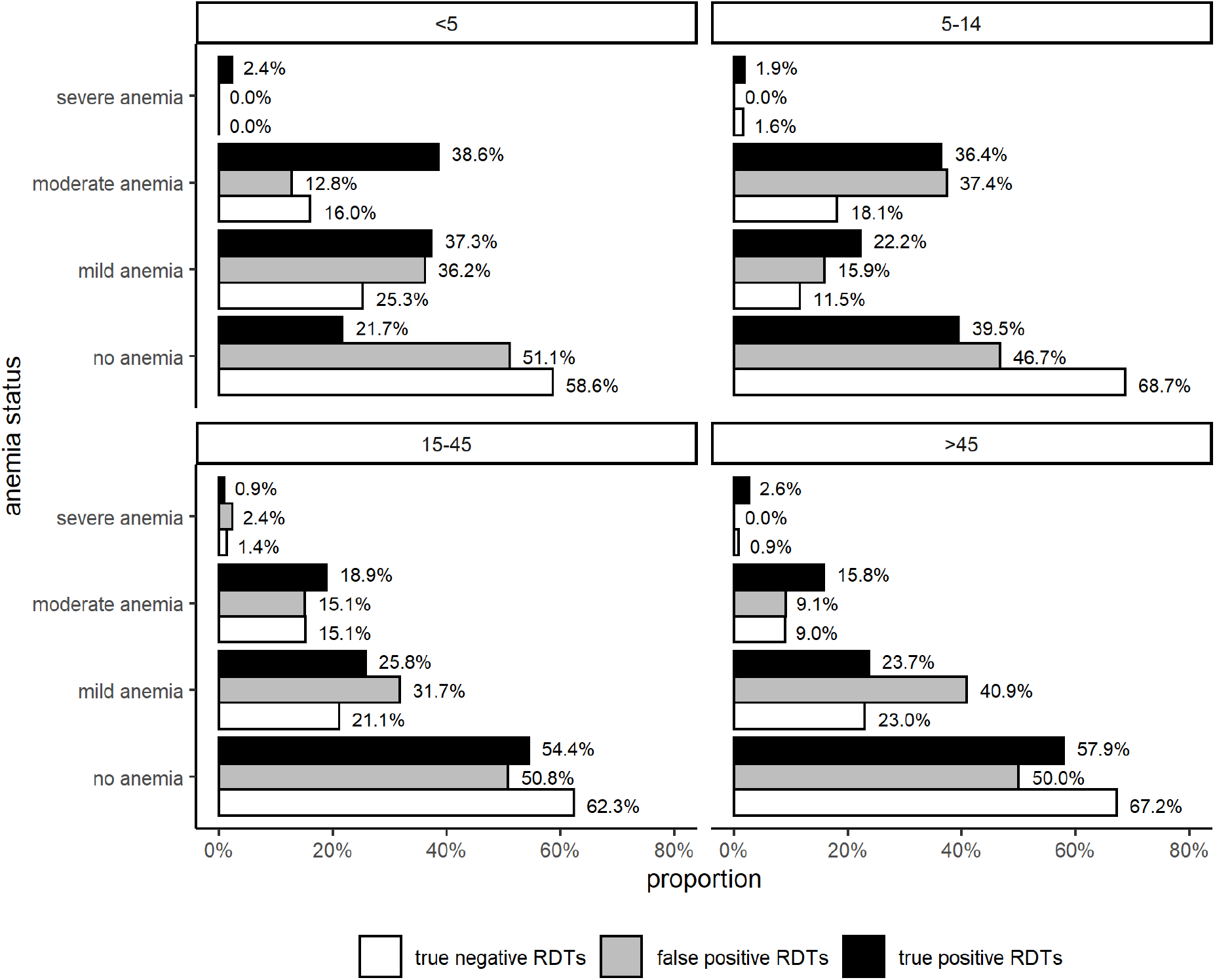
Anaemia status compared between true positive, false positive, and true negative RDT outcomes and stratified by age.

## Discussion

Malaria control programs rely on continuous and systematic collection of surveillance data for decision making and resource allocation ^34^. A critical measure that closely reflects malaria transmission intensity is the parasite rate, the proportion of the population found to carry parasites in their peripheral blood ^35^. RDTs, more specifically PfHRP2-based RDTs, are the most widely used test to measure parasite rates in endemic countries and are a cornerstone of malaria control. However, here we have identified diagnostic performance issues, particularly related to limited specificity. Therefore, malaria surveillance depending solely on RDTs needs well integrated quality control procedures assessing the impact of reduced sensitivity and specificity of the tests used in malaria control programs. In this report, we present an efficient approach to assess the performance of field-deployed RDTs used for malaria surveillance on large scale based on molecular analysis of NA retained on the RDTs.

In 4.7% (84/1800) of the negative RDTs we found *Plasmodium* spp. NA and identified these as FN-RDT. The low proportion of FN-RDT can be explained by the low parasite densities of these asymptomatic individuals (geometric mean of 5.4 Pf/μL) and the low amount of blood (one drop corresponds to approximately 5 μL) used as starting material for the molecular analysis. This is a limitation of our approach. Therefore, the true proportion of FN-RDTs in a high prevalence setting such as Bioko Island is likely to be higher.

We identified *Pf* isolates with potential *pfhrp3* deletions but not a single isolate with a confirmed *pfhrp2* deletion. Given the overall high frequency of polyclonal *Pf* infections in this setting (63% by *pfmsp1/pfmsp2* genotyping), we assumed that if *Pf* carrying *pfhrp2* deletions exist, then they would be most likely masked by co-infecting *Pf* isolates without *pfhrp2* gene deletions. Of all the samples included for final analysis, 30.6% had an increased ΔCq value for *pfhrp2* and 50.0% for *pfhrp3* amplification, indicating for the first time that there are likely *Pf* strains circulating on Bioko Island carrying deletions in their *pfhrp2* and/or *pfhrp3* genes. So far, one report described *Pf* strains carrying *pfhrp2* and *pfhrp3* deletions in blood samples collected on the continental region of Equatorial Guinea ^36^. However, since the travel activity between Bioko Island and the mainland of Equatorial Guinea is high, it can be assumed that parasite strains are exchanged frequently between these locations ^37^. Most importantly, blood samples with *Pf* clones indicative for masked *pfhrp2* and *pfhrp3* gene deletions were recorded as PfHRP2 positive by RDT. Likely, the co-circulating *Pf* clones compensate for the lack of PfHRP2 expression resulting in RDT positive testing.

Our data support the notion that in malaria medium to high transmission settings, where polyclonal *Pf* infections are common, only assays with the ability to identify masked *pfhrp2* and/or *pfhrp3* gene-deleted parasites should be used ^38^. Importantly, to avoid false reporting of *pfhrp2* and/or *pfhrp3* gene deletions, we used a robust and multi-layered approach by which only samples with a pre-defined parasite density, successful amplification of the assays’ internal control, and additional, independent amplification of either *pfmsp1* or *pfmsp2*, were included into the analysis.

In our study, we discovered a significant proportion of FP-RDTs that were declared as malaria positive in the field. Our findings are not unique to Bioko Island. In a study conducted in Tanzania, 22% of malaria positive RDTs were negative by molecular analysis for *Pf* ^39^. A study performed in Guinea-Bissau reported 26% FP-RDTs ^40^, and in Western Kenya, approximately one-third of positive RDTs were negative by molecular detection methods for *Pf* ^41^. With the wider access to novel “ultra-sensitive” RDTs, which are detecting lower concentrations of the PfHRP2 antigens, the problem of FP-RDT results is expected to become even greater, as already shown in a recently published study ^42^.

The PfHRP2 antigen (97.4%) was much more often detected among FP-RDTs than the pLDH antigen (11.2%). We were able to associate the false positivity of PfHRP2-RDTs with the recent use of antimalarial drugs. It has been well established that antimalarial treatment leads to false positive PfHRP2-RDT results because the PfHRP2 antigen persists in the blood days to weeks after parasite clearance ^22, 43–46^.

Remarkably, we found not only an association between the recent use of antimalarial drugs and FP-RDTs, but also an indirect association with accessibility to antimalarial drugs. A higher socio-economic status and living in the urban part of the Island directs towards increased accessibility to antimalarial drugs.

The impact of FP-RDTs differs greatly depending on the setting in which RDTs are deployed. In clinical settings, FP-RDTs might be less common, but the consequences are more serious since unnecessary prescription of antimalarials might increase the risk of overlooking other, life-threatening diseases causing fever or inducing side-effects caused by the antimalarial drugs ^47^. However, if RDTs are used for epidemiological surveys, a high proportion of FP-RDTs due to PfHRP2 antigen persistence might lead to an overestimation of the malaria prevalence in regions with good access to antimalarial treatment (urban regions) or in populations which are more likely to receive antimalarial treatments (higher socio-economic status). One striking example shown here is that the use of RDTs to assess the relationship between asymptomatic malaria infection and anemia status in children < 5 years will lead to an underestimation of the severe consequence of asymptomatic malaria infections on hemoglobin levels.

The benefits and the challenges that come with large-scale deployment of molecular techniques in malaria endemic regions have been discussed elsewhere ^48^. And now, the COVID-19 pandemic has renewed the discussion regarding the importance of molecular testing as part of public health systems across Africa ^49^ and will likely accelerate efforts to integrate molecular tools for larger scale genomic surveillance of malaria into control programs.

## Conclusion

Malaria surveillance programs based on RDT assessments of malaria prevalence should be strengthened by the integration of molecular epidemiological data in the same setting. These data will serve as an early warning system for (i) spread of *Pf* strains evading widely used diagnostic tests, (ii) understanding overuse of malaria drugs, (iii) help with identifying fever causing diseases beyond malaria, and (iv) help to clarify the burden of asymptomatic malaria as a cause of severe to moderate anemia, particularly in children < 5 years.

## Abbreviations

Pf: (*Plasmodium falciparum*)
RDT: (rapid diagnostic test)
PfHRP2/3: (histidine rich protein 2/3)
ENAR: (Extraction of nucleic acids from RDTs)
MIS: (Malaria Indicator Survey)
RT-qPCR: (reverse transcription quantitative polymerase chain reaction)
Cq: (quantification cycle)
sSA: (sub-Saharan Africa)
ACT: (artemisinin combination therapy)
SES: (socio-economic status)
NA: (nucleic acid)

## Acknowledgments

The authors would like to thank all MIS participants for their contribution and the BIMCP staff for their commitment and support during sample collection. We would like to thank Christin Gumpp, Christian Scheurer and Sergio Wittlin from the Swiss TPH Malaria Drug Discovery Group for their help with cultivating PfNF54, PfDD2 and PfHB3 parasites, whose DNA was used as controls for the qHRP2/3-del assay. We would like to acknowledge Amanda Ross for her support and guidance with the statistical analysis used in this manuscript.

## Author contributions

Conceptualization: SH, CD, TS. Data curation and validation: SH, TS, OTD. Formal analysis and visualization: SH. Funding acquisition: CD, MT, GAG, WPP. Investigation: OTD, GAG, WPP, MOA, CAG. Methodology: SH, CAY, EAG, JPD, KB. Resources: MM, EN, OTD, GAG, WPP, CAG. Project administration and supervision: CD, TS. Writing original draft: SH, TS, CD. All authors reviewed the manuscript.

## Supplementary Materials

**Supplementary figure 1.**
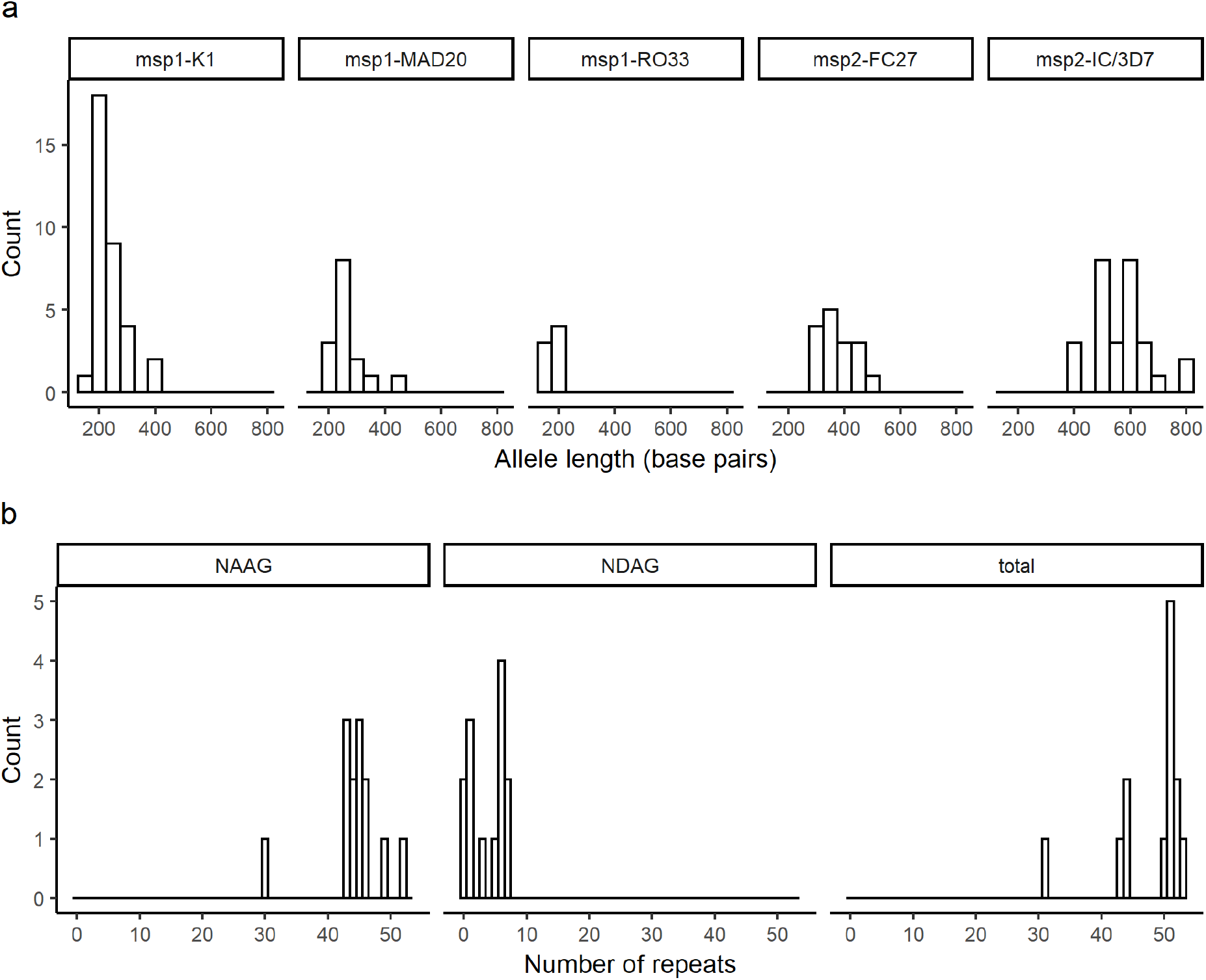
(a) Genetic diversity of *Pf* determined by size polymorphism of *pfmsp1* and *pfmsp2*. (b) Genetic diversity of *P. malariae* determined by the number of NAAG and NDAG repeats in the *pmcsp* gene.

**Supplementary figure 2.**
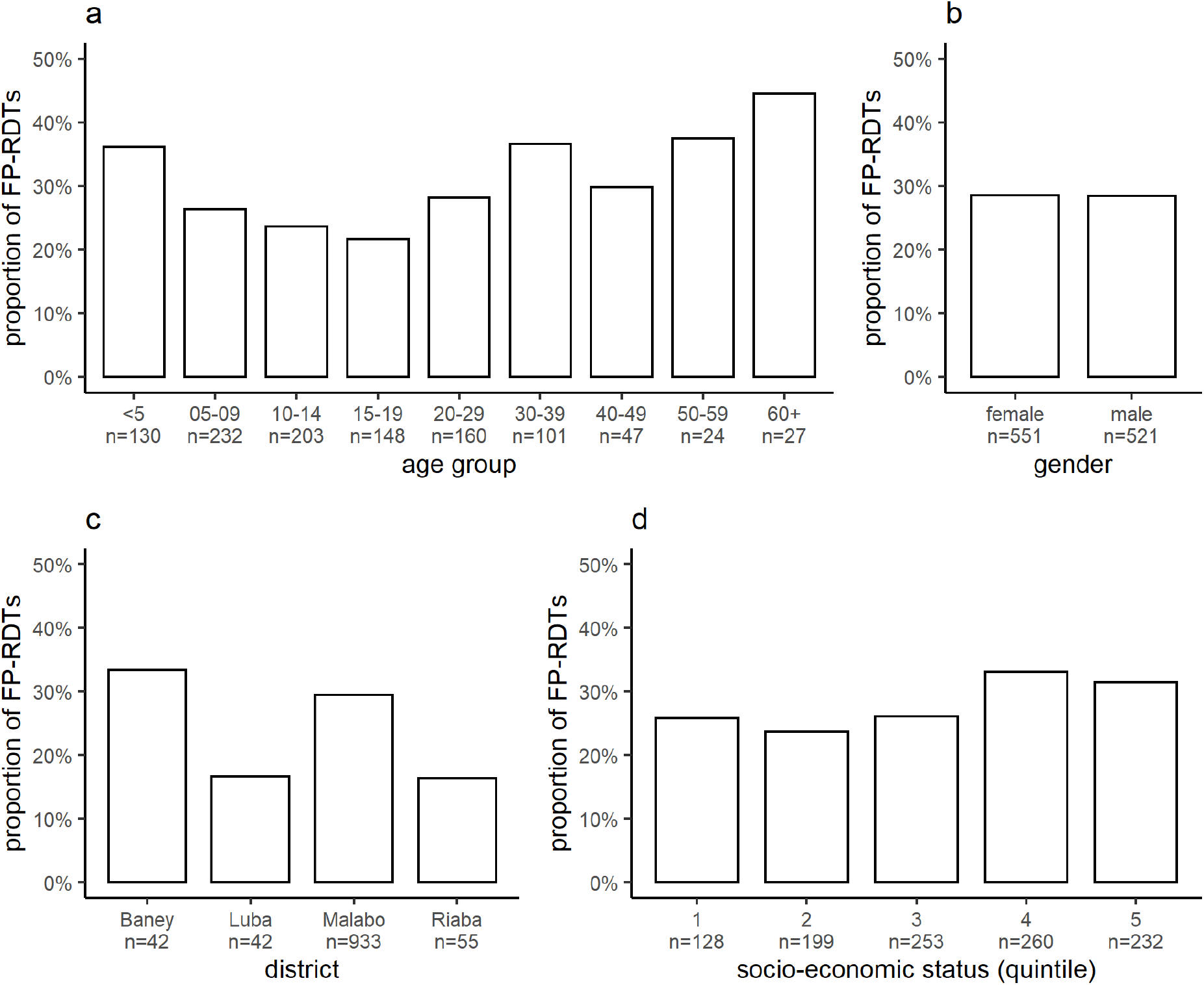
FP-RDTs as a proportion of all positive RDTs stratified by age (a), gender (b), district (c) and socio-economic status (d). Quintile 1 refers to the lowest, while quintile 5 to the highest socio-economic status.

**Supplementary table 1:**
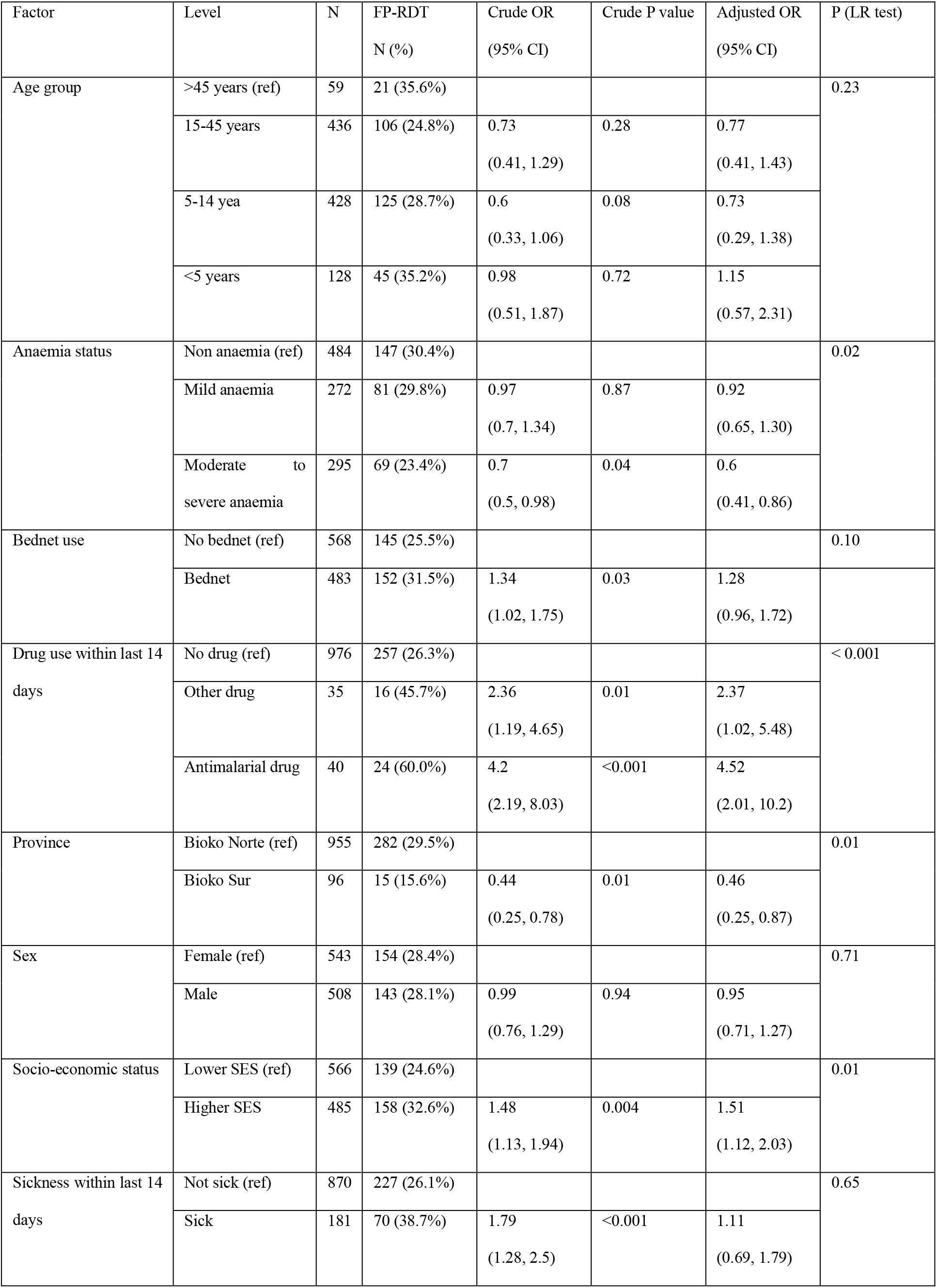
Multivariable logistic regression analysis of risk factors associated with FP-RDTs using community as a random effect. Crude and adjusted odds ratios and their respective 95% confidence intervals were calculated based on comparison between FP-RDT and TP-RDT.

